# Exploring the kinetics and mechanism of phase separation in ternary lipid mixtures containing APP C99 using atomistic vs coarse-grained MD simulations

**DOI:** 10.1101/2024.12.31.630974

**Authors:** George A. Pantelopulos, Sangram Prusty, Asanga Bandara, John E. Straub

## Abstract

The phase separation of lipid bilayers, composed of mixtures of saturated and unsaturated lipids and cholesterol, is a topic of fundamental importance in membrane biophysics and cell biology. The formation of lipid domains, including liquid-disordered domains enriched in unsaturated lipids and liquid-ordered domains enriched in saturated lipids and cholesterol is believed to be essential to the function of many membrane proteins. Experiment, theory, and simulation have been used to develop a general understanding of the thermodynamic driving forces underlying phase separation in ternary and quaternary lipid mixtures. However, the kinetics of early events in lipid phase separation in the presence of transmembrane proteins remain relatively understudied. Using large-scale all-atom and coarse-grained simulations, we explore the kinetics and phase separation of ternary lipid mixtures of saturated lipid, unsaturated lipid, and cholesterol. Order parameters employed in the Cahn-Hilliard theory provide insight into the kinetics and mechanism of lipid phase separation. We observe three distinct time regimes in the phase separation process: a shorter time exponential phase followed by a power law phase followed by a longer time plateau phase. Comparison of lipid, protein and lipid-protein dynamics between all-atom and coarse-grained models identifies both quantitative and qualitative differences and similarities in the phase separation kinetics. Moreover, timescaling of dynamics of AA and CG simulation yields a similar kinetic mechanism of phase separation. The findings of this study elucidate fundamental aspects of membrane biophysics and the ongoing efforts to define the role of lipid rafts in the structure and function of cellular membrane.

## 1 Introduction

Lipid mixtures composed of saturated and unsaturated lipids in conjunction with cholesterol in appropriate proportion, are observed to undergo phase separation. ^1–4^ Cholesterol has a higher affinity for associating with saturated lipids compared to unsaturated lipids. ^5,6^ Domains formed by the association of predominantly saturated lipids and cholesterol adopt a liquid-ordered (L*_o_*) phase when the cholesterol mole fraction exceeds 0.2 (20 mol%). ^7–10^ The L*_o_* domains are characterized by a smaller area per lipid, increased membrane thickness, and more ordered lipid tails. ^11,12^ The higher density observed in liquid ordered domains leads to hexagonal packing of lipid tails at lower mol% cholesterol ^9,13^ and the formation of cholesteric gels at very high concentrations characteristic of ocular membranes. ^14^

“Lipid rafts” are evidenced to be principally defined by the L*_o_* phase, and have been hypothesized to play a functional role in cellular metabolism. ^15,16^ It has been conjectured that certain membrane proteins are recruited to raft domains, at times facilitated by post-translational lipidation, and colocalized with related proteins or substrates in a way that facilitates function. ^17–19^ One example is found in the amyloid cascade associated with Alzheimer’s disease. Evidence suggests that the enzymes β-secretase and γ-secretase are colocalized in raft domains and carry out correlated cleavage of the Amyloid Precursor Protein (APP) substrate, C99. ^20–23^

### 1.1 Thermodynamics of lipid phase separation

The phenomenon of lipid phase separation and raft domain formation has received a great deal of attention in experimental and theoretical studies. ^1,24–29^ The seminal work of Keller and Veatch ^30^ established a minimal experimental model exhibiting binary phase separation and the formation of L*_o_* lipid domains involves a ternary mixture including saturated lipid, unsaturated lipid, and cholesterol. In addition, simple statistical models of binary phase separation, such as Ising models^31,32^ and Flory-Huggins models, ^24,33^ have been used to capture key features of the observed process of phase separation. However, phase separation may also be observed in more complex mixtures, including sphingophospholipid, sphingomyelin, glycolipids GM1 and GM3, and even proteins. ^34–41^

Computer simulations of lipid phase separation employing coarse-grained (CG) and allatom (AA) models have been used to study the nature of L*_d_* and L*_o_* phases. Spontaneous phase separation from ternary lipid mixtures was first observed in computational studies employing CG models. ^42,43^ Early studies of L*_d_* and L*_o_* states using AA models consisted of single phase systems or systems that were initially phase separated. ^8^ However, more recent studies using AA models have observed initial stages of phase separation from an initially mixed state. ^44^

Theoretical arguments, based on Flory-Huggins models of binary phase separation, and systematic computational studies, employing CG models, have demonstrated that systems on the order of a thousand lipids are required to create thermodynamically stable phase separated states. ^33^ The mixed state is favored by entropy, through the entropy of mixing. The phase separated state is favored by enthalpy, where saturated lipids engage in favorable van der Waals interactions following dissociation from unsaturated lipids and association with cholesterol, transitioning from the L*_d_* phase to the L*_o_* phase. Additional contributions to stabilizing entropic changes in lipid tail^45^ and lipid-water conformations may also contribute to stabilizing the phase-separated state. ^46^ However, there is a free energy cost to create an interface between the L*_o_* and L*_d_* domains. The system size must be large enough for the enthalpic gain of forming the L*_o_* domains can compensate for the cost of demixing and formation of the domain interface.^47^

### 1.2 Kinetics of lipid phase separation

While the fundamental thermodynamic driving forces responsible for lipid phase separation into L*_d_* and L*_o_* domains is well established, relatively little is known about the early stages of lipid phase separation kinetics. The times scales, presumed to be tens of milliseconds, and length scales, estimated to be on the order of a tenth of a micron, for phase separation into L*_o_* domains pose challenges for experiment.^48,49^ These length scales are at the limit of super-resolution microscopy. Chromophores like amide bands in sphingophospholipids are sensitive to hydrogen bonding with solvent or protein, but they show limited sensitivity to the membrane phase, resulting in the absence of reliable spectroscopic indicators for membrane phase detection.^50^

Relatively little insight into the kinetics and mechanism of lipid phase separation has been provided by simulation using CG or AA models. The simulation study of Bennett et. al., ^51^ observed phase separation in a ternary lipid mixture involving cholesterol on a time scale of tens of microseconds. While the study included a non-equilibrium simulation of phase separation, the primary focus of the work was thermodynamics. They observed that the CG model was able to capture the enthalpic stabilization on forming the L*_o_* phase, but failed to capture the loss of entropy associated with lipid tail ordering. However, the cost in free energy associated with establishing the domain interface, found to be critical in other studies, was not discussed. Free energies as a function of lipid mixing have been determined for the MARTINI2 coarse-grained model. ^52^ A barrier between miscible and phase separated states of approximately 1 kcal/mol was observed, similar to estimates based on lipid domain line tensions^33,47^.

A significant number of studies have employed free energy functional approaches to explore the kinetics of phase separation in lipid mixtures. ^53–58^ Those studies have often represented the state of the system in terms of a space- and time-dependent field variable representing the local lipid composition. Many important observations have been made regarding the nature of the kinetics of lipid phase separation, including the roles of nucleation and growth, domain ripening, coalescence, and budding, ^54^ as well as the role of hydrodynamics in the membrane and solvent.^59^

Of particular importance are considerations of the role of line tension. Though the line tension on the nanoscopic scale has not been estimated experimentally, it was noted that lower line tensions would be consistent with the formation of lipid raft microdomains that exist in a quasi-equilibrium with larger scale domains. ^59–62^ In the event that the line tension is larger, lipid rafts were proposed to form through a process involving the wetting of proteins by saturated lipids and cholesterol. The coalescence of similarly lipid-wetted proteins could lead to larger scale formation of raft domains containing multiple proteins. ^63–65^ Alternatively, it has been proposed that nanoscopic raft domains could be formed through the process of exocytosis or endocytosis. ^59^

A notable recent study of the kinetics of lipid phase separation using AA molecular dynamics included simulation bilayers formed by 1,2-dipalmitoyl-*sn*-glycero-3-phosphocholine:1,2-dioleoyl-*sn*-glycero-3-phosphocholine (DPPC:DOPC) and DPPC:DOPC:cholesterol lipid mixtures over a range of temperature from 270K to 310K. ^44^ Over the course of ten microsecond-long simulations, the phase separation process was observed to continue, indicating that the time scale for phase separation was on the order of tens of microseconds. In ternary mixtures, phase separation into gel-L*_o_* phases was observed at 270K and L*_o_*-L*_d_* phases at 280K and 290K, while a single L*_d_* phase was observed at 310K. However, no particular insight into the mechanism of phase separation was noted.

In this work, we perform large-scale molecular dynamics simulations of a pair of C99 protein conjugates in a ternary lipid mixture consisting of 37 mol% saturated lipid, DPPC, 37 mol% unsaturated lipid, 1,2-dilinoleoyl-*sn*-glycero-3-phosphocholine (DUPC), and 26 mol% cholesterol (CHOL) at atomistic and coarse-grained resolution. Our study explores the kinetic mechanism of lipid phase separation in the presence of transmembrane proteins and provides a comparative analysis between AA and CG models. Notably, we find that after appropriate timescaling, the kinetic mechanisms of phase separation are similar in both models, despite some differences in the early stages of phase separation. Furthermore, we assess the monomeric and dimeric states as well as the localization of the protein in both models. Overall, our findings provide insight into the kinetic mechanism of lipid phase separation and microdomain formation in ternary lipid mixtures in presence of APP C99 protein, along with highlighting the importance of timescaling when comparing dynamics at different levels of resolution.

## 2 Methods

### 2.1 All-atom simulation on ANTON2

We performed a single simulation of a 496,452-particle system containing a lipid bilayer using the NPT ensemble for 22 µs of sampling. The system was composed of two 40-residue transmembrane proteins, 640 DPPC lipids, 640 DUPC lipids, and 480 CHOL in a symmetric lipid bilayer. The membrane and protein system was solvated in 96800 TIP3P waters and 139 Na^+^ and 149 Cl*^−^* ions. The initial conditions of the system consisted of a random lateral mixture of lipids and two copies of APP-C99_16*−*55_ proteins (C99) separated by a distance equivalent to one lipid solvation shell apart with the juxtamembrane domains of each C99 structured into an alpha helix and oriented toward each other based on PDB:2LLM. ^66^ The CHARMM36^67^ force field family of parameters were used for all components of the system. The initial conditions were prepared and equilibrated using CHARMM-GUI^68^ protocols.

The production simulation was run on ANTON2 ASIC19^69^ in an NPT ensemble using a 2.4 fs time step and a RESPA^70^ scheme with the default bonded interval of 1-time step, far non-bonded interval of 3-time steps, and non-bonded near interval of 1-time step. The pressure was maintained at 1 bar and a compressibility of 4.5 × 10*^−^*^5^ bar*^−^*^1^ with an MTK barostat^71^ semi-isotropic pressure coupling scheme. The temperature was coupled to a Nośe-Hoover thermostat^72,73^ at reference temperature of 299 K, which was required to ultimately achieve an average simulation temperature of 295 K. Thermostat and barostats were each coupled at 24 ps intervals. Long-range interactions were computed using the u-series method with default parameters. Center of mass motion removal was applied to prevent incidental drift. Thermodynamic variables were saved every 2.4 ps and coordinates were saved every 240 ps. The Anton2 trajectory reached 18 µs. The trajectory was extended to 22 µs in GROMACS using the leap-frog integrator with a 2 ps time step, and the semiisotropic Parrinello-Rahman barostat for pressure coupling ^74^ and Nośe-Hoover thermostat for temperature coupling.

### 2.2 MARTINI2 and MARTINI3 simulations on CPUs+GPU

We performed five 12 µs simulations of 43,846-particle systems of identical membrane composition to the all-atom systems described in the previous subsection with the MARTINI2^75,76^ and MARTINI3^77,78^ coarse-grained force fields. Due to the MARTINI 4-to-1 all-atom to coarse-grained mapping of water, we represent the solvent with 24,200 water particles, 10% of which are “anti-freeze” water to prevent spontaneous freezing in MARTINI2 simulation, and 139 Na^+^ and 145 Cl*^−^* ions. The same initial configuration, spacing, and relative orientations of the two C99_16*−*55_ as in the all-atoms systems were used in all coarse-grained simulations. The initial coordinates of each of the 5 replicates systems in MARTINI2 and MARTINI3 used the same initial lipid and protein positions by producing initial coordinates from 5 well-mixed representations of these systems in MolPainter. ^79^ Each system was minimized and then annealed with a 10 fs time step over 2 ns from 200 to 295 K and equilibrated for 1 ns at 295 K using the Berendsen thermostat and semi-isotropic barostat at 1 bar and 3×10*^−^*^4^ bar*^−^*^1^ compressibility using the same non-bonded pair interaction calculation scheme as in production simulations.

Production simulations were run on the Boston University SCC cluster with 7 Intel Xeon E5-2680v4 CPUs and 1 P100 GPU for each replica using a 25 fs time step and a leap-frog integration scheme using GROMACS 2021.5. ^80^ The pressure was maintained at 1 bar with a Parrinello-Rahman barostat semi-isotropic pressure coupling scheme coupled at a 12 ps interval and a compressibility of 3 × 10*^−^*^4^ bar*^−^*^1^.^74^ The temperature was maintained at 295 K with the Bussi velocity-rescale thermostat coupled at a 1 ps interval for separate groups of the membrane, including protein and the solvent.^81^ Lennard-Jones pair interactions were computed and shifted to zero at 1.1 nm. Electrostatic pair interactions were computed using the reaction-field scheme with a 1.1 nm cutoff and a dielectric of ɛ = 15.

## 3 Results and Discussion

### Both AA and CG models display spontaneous phase separation to form L*_o_* and L*_d_* domains

Instantaneous configurations depicting the spatiotemporal evolution of lipid phase separation are presented for the AA model simulation in Figure 1. Over the course of the 22 µs simulation, the lipid phase separation spontaneously proceeds from a miscible state to small islands like L*_o_* (bluish) and L*_d_* (reddish) aggregates, which eventually merge to form a complete phase-separated L*_o_* and L*_d_* domains. The two C99 proteins (yellow) are found to be localized at the domain interface and primarily solvated by unsaturated DUPC lipids. Moreover, with the progress of the simulation the APP C99 protein is observed predominantly in monomeric state in the AA simulation.

**Figure 1:**
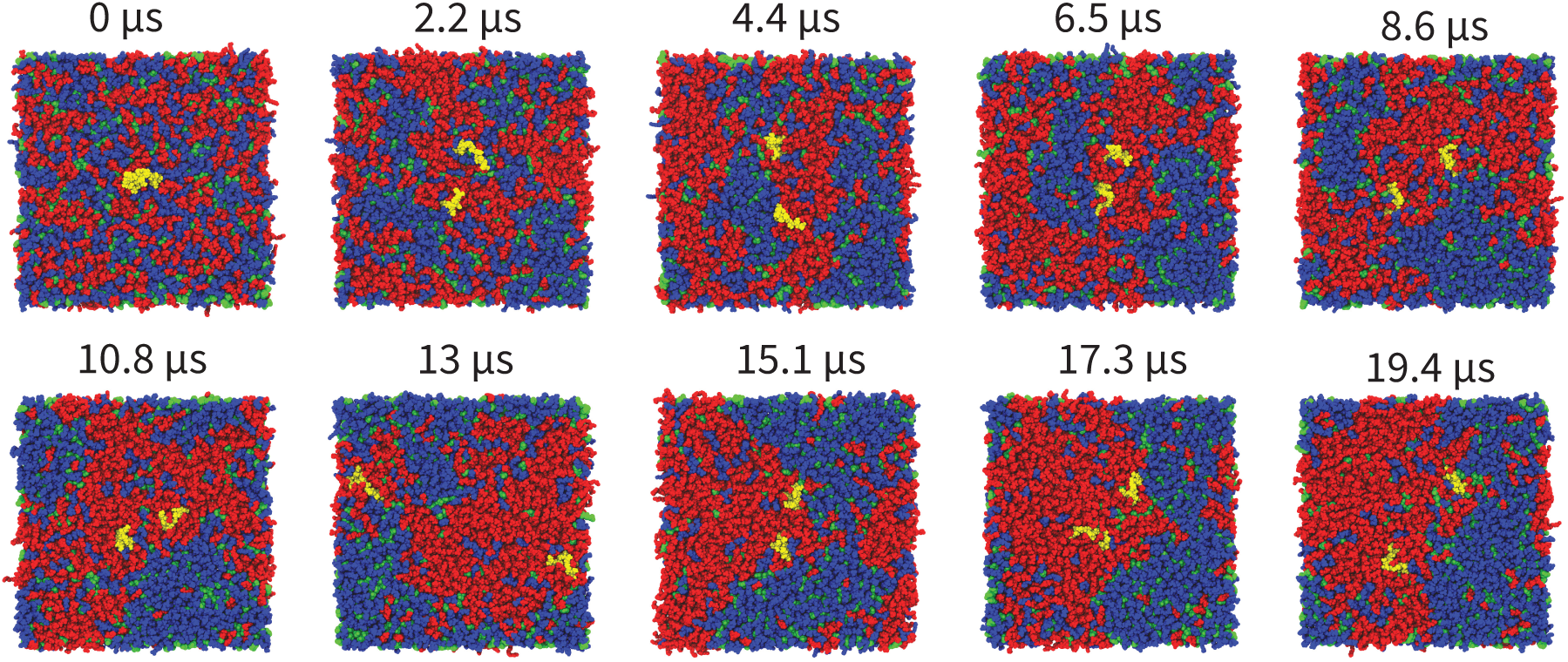
Ten instantaneous configurations depicting the process of lipid phase separation of DPPC (blue), DUPC (red), and cholesterol (green) into L_o_ (bluish) and L*_d_* (reddish) regions. The C99 proteins (yellow) are found to be positioned predominantly within the L*_d_* phase at the domain interface.

A similar spontaneous phase separation process is also observed for CG simulation as shown in Figure S1. However, a significant difference is observed in the protein localization and oligomeric state between the AA and CG simulations. In the MARTINI2 simulations, the protein is predominantly localized at the domain interface. In contrast, in MARTINI3 simulations, the protein shifts toward the L*_d_* domain, suggesting difference in lipid-protein interactions between the two MARTINI models. Furthermore, in contrast to the AA simulations where the protein predominantly exists as a monomer, both MARTINI2 and MARTINI3 simulations observe the protein in a dimeric state. This difference in oligomeric state highlights the differences in protein-protein interaction between AA (CHARMM36) and CG (MARTINI2 and MARTINI3) models.

### Timescaling of dynamics in AA and CG simulations yields similar time evolution

A useful quantitative measure of the progression of phase separation is the mixing entropy S_mix_(t) as shown in Figure 2 for the AA and CG dynamics. Following our early studies of phase separation in ternary lipid bilayers, ^33^ the mixing entropy was defined in terms of the probability of contacts formed by the same lipid type (p_1_) or different lipid type (p_2_)

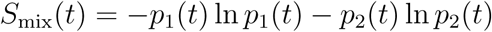

**Figure 2:**
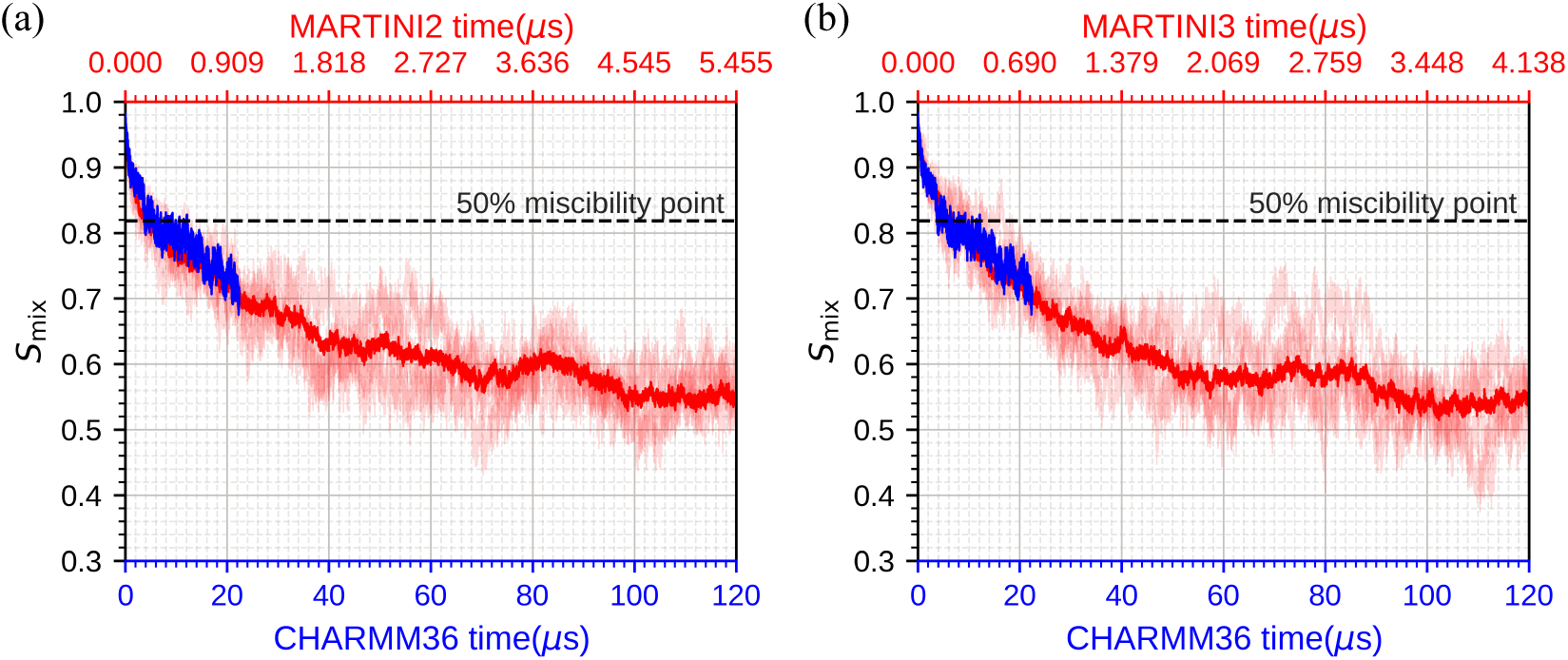
The time evolution of the mixing entropy S_mix_ for the AA CHARMM36 (blue) and CG (red)(a) MARTINI2 and (b) MARTINI3 simulations. The time axes depicts the evolution of the time in each simulation where the CHARMM time is approximately 22 times longer than the MARTINI2 time and 29 times longer than the MARTINI3 time at a similar degree of phase separation.

The local lipid contacts were computed using nearest neighbors determined via Voronoi tessellations. This method provides advantages over definitions of mixing entropies based on coarse-grained measurements of lipid composition within a space-fixed grid, the resolution of which influences the estimate of the mixing entropy. Atoms closest to the lipid leaflet mid-plane, two from lipids and one from cholesterol, were selected within each leaflet in each frame, from which a Voronoi tessellation was performed.

Figure 2a demonstrates the comparison of time evolution of S_mix_ in CHARMM36 (AA) and MARTINI2 (CG) simulations. The time axes of MARTINI2 (red) have been scaled with respect to CHARMM36 (blue) timescale so as to provide best fit of the two time series. Similarly Figure 2b provides the comparison between CHARMM36 (AA) and MARTINI3 (CG) simulations. The scaling factor has been calculated using a χ^2^ analysis as demonstrated in Figure S2. For CHARMM36 and MARTINI2 simulations the scaling factor has been calculated to be 22 and for CHARMM36 and MARTINI3 simulations it has been calculated to be 29.

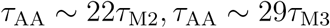

The miscible state is represented by S_mix_ ≈ 1 and a completely phase separated state by S_mix_ ≈ 0 for a bulk system. For a finite size system showing some impurities within a given lipid phase, the value of S_mix_ for the phase separated state will be greater than zero. As is evident from the Figure 2 the process of phase separation is incomplete in both the AA and CG simulations. Moreover, the time-dependence of the S_mix_(t) is qualitatively similar in the AA and CG simulations after time scaling. The 50% miscibility point^9^ is reached in approximately 5µs of CHARMM time, 230ns of MARTINI2 time and 170ns of MARTINI3 time.

Another useful measure of the progress of phase separation is the size of the maximum cluster representing a L_o_ cluster of DPPC and cholesterol. This measure is expected to approach a constant value representing the phase separated state. Figure 3 depicts the maximum cluster size as a function of time. The time axes has been scaled based on the scaling factor obtained for S_mix_. Figure 3a and Figure 3b illustrate the qualitative similarity observed between the evolution of maximum cluster size in AA (CHARMM36) and CG (MARTINI2 and MARTINI3) dynamics upon time scaling, with the CG dynamics approaching convergence. However, greater difference between the two time series is evident on the log-log plot as shown in Figure 3c and Figure 3d. Even after the time scaling, the CG dynamics is observed to have a faster early stage of growth of the maximum cluster size, while the AA dynamics presents a noticeable lag phase.

**Figure 3:**
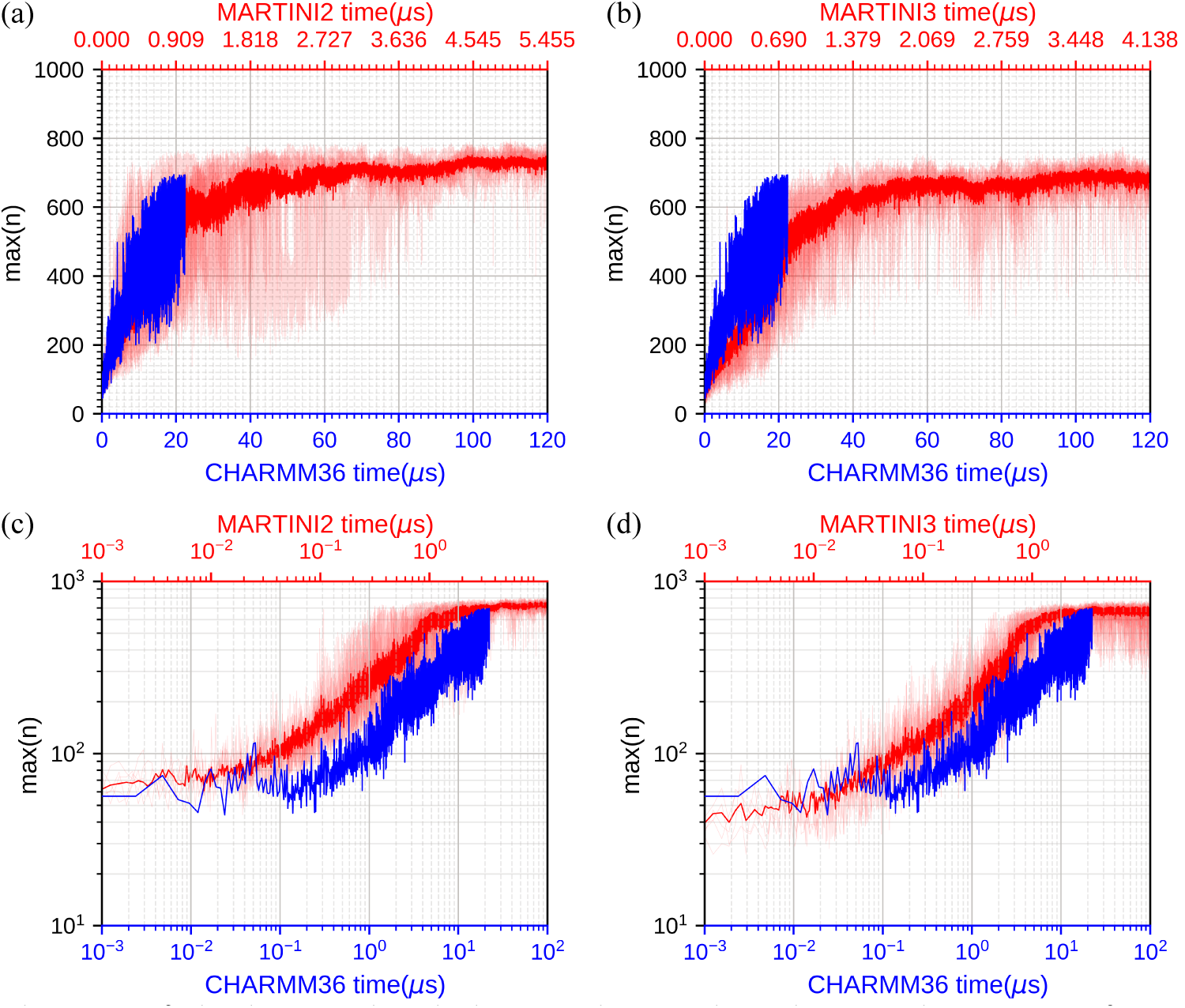
The size of the largest lipid cluster observed in the simulation as a function of time for AA and CG trajectories. The time axis is scaled using the same scaling factor as that of S_mix_. The top panel demonstrates the comparison between the time evolution of maximum cluster size between CHARMM36 and (a) MARTINI2, (b) MARTINI3 simulations. The bottom panel demonstrates the time evolution on a log-log scale for the top panel comparisons. Following time scaling, general agreement between the AA and CG results is observed (top) however, on a log-log scale (bottom), a greater rate of initial growth of the maximum cluster size is observed in the CG simulation.

The size of the maximum cluster increases as a power-law after 20ns in MARTINI2 timescale till the convergence point:

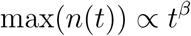

where fitting after scaling to MARTINI2 timescale leads to exponents of β_AA_ = 0.52, β_M2_ = 0.54 and β_M3_ = 0.55. These exponents are indicative of a diffusion-limited coalescence being the dominant mechanism for domain growth over the course of the simulation. A coalescence mechanism of domain growth is expected as (1) the composition of lipids forming L*_o_* and L*_d_* domains is roughly equal, and thus their domain areas are approximately equal, and (2) the time scale for evaporation of lipids from domains is large compared with the simulation time scale.

### Origin of timescaling in terms of higher diffusion and sticking probabilities

To understand the origin of the empirical scaling factor between the time evolution of AA and CG simulation, we develop a simple model for the nucleation and growth of a growing cluster. Let i be the number of monomers in a cluster n. The reversible reaction for addition of another monomer to the cluster n can be written as:

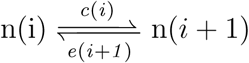

where c(i) is the rate of condensation of a monomer onto a cluster of i monomers and e(i+1) is the rate of evaporation of a monomer from a cluster of i+1 monomers. If condensation occurs through diffusion controlled encounter, the rate of addition of monomers is approximately

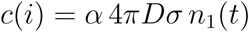

where D = D*_i_* +D_1_ is the relative rate of diffusion of the cluster and monomer and σ = r*_i_* +r_1_ is the encounter radius for contact between the cluster and monomer. The parameter α is a sticking probability that represents the fraction 0 ≤ α ≤ 1 of collisions that lead to the addition of a monomer to the cluster. Therefore, the ratio of the rate of growth of a cluster in AA and CG simulation can be given by:

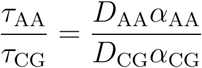

Thus, the emergence of scaling factor can be attributed to differences in the diffusion coefficients and the sticking probability for the AA and CG dynamics.

In order to get a quantitative insight into the magnitude of contribution of diffusion coefficient to that of the scaling factor we have compared the lateral diffusion constant of DPPC, DUPC, and Cholesterol obtained from AA and CG simulations in this section. We have calculated the lateral diffusion constant from the slope of the average mean square displacement vs time plots (Figure S3), excluding the first and last 10% of the trajectory. For the CG simulations, we used five replicas each of MARTINI2 and MARTINI3 simulations to calculate the average mean square displacement.

Figure 4 shows a comparison of the diffusion constant for each of the lipids involved in the AA and CG simulations. We observe an enhancement in the diffusion constant for MARTINI(CG) simulations relative to AA simulations. Table 1 presents the ratio of the diffusion coefficient of CG dynamics to that of AA dynamics for DPPC, DUPC, and Cholesterol. We note that the highest diffusion rates are observed for Cholesterol. Considering the highest ratio of diffusion constant of 8.4 and 12.6 for MARTINI2 and MARTINI3 simulations respectively, we obtain a missing factor of ∼ 2.6 and ∼ 2.3 for MARTINI2 and MARTINI3 simulations. These factors can be attributed to the sticking probability ratios as per the model which we developed above.

**Figure 4:**
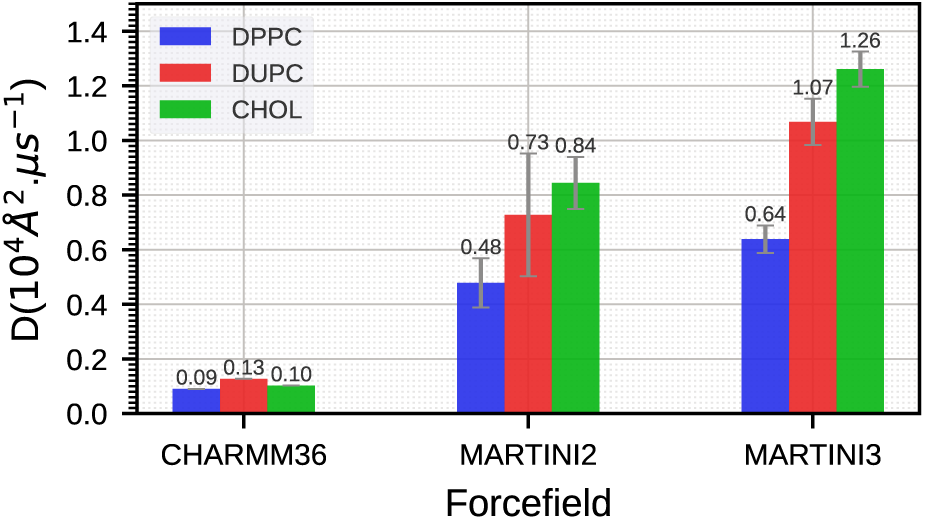
Mean diffusion constant for DPPC, DUPC and Cholesterol as obtained from a single AA (CHARMM36) trajectory and 5 replicas of CG (MARTINI2, MARTINI3) trajectories. MARTINI3 exhibits the highest diffusion constant for all lipids, which can be attributed to the fastest rate of phase separation.

**Table 1:** Ratio of Diffusion constant for lipids in CG simulation to AA simulations.

| Lipid | $D_{M2}/D_{AA}$ | $D_{M3}/D_{AA}$ |
| --- | --- | --- |
| DPPC | 5.3 | 7.1 |
| DUPC | 5.6 | 8.2 |
| CHOL | 8.4 | 12.6 |

### Three-stage mechanism of phase separation

Several studies have utilized the Cahn-Hilliard theory extensively in order to understand phase separation kinetics. However, most of the studies have employed hydrodynamic approaches that are only applicable to larger lengths and timescales, and thus cannot capture the microscopic picture. Bagchi and co-workers ^82^ were the first to apply the concepts from the Cahn-Hilliard theory to MD simulations in order to understand the microscopic origin of the different stages involved in the kinetics of spinodal decomposition in binary mixtures. Building upon Bagchi’s work, we developed the Cahn-Hilliard local composition order parameter(ϕ(**r**, t)) to study the different stages involved in the kinetics of lipid phase separation in ternary lipid mixtures. It is defined as

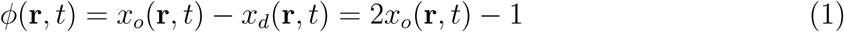

where x*_o_*(**r**, t) and x*_d_*(**r**, t) are the mole fractions of saturated and unsaturated lipids. For a miscible state, we expect ϕ ≈ 0, while for liquid ordered or liquid disordered regions we expect ϕ ≈ 1 and ϕ ≈ −1, respectively. During the process of phase separation, the probability distribution p(ϕ) reflecting the range of observed values of ϕ(**r**, t) at a given time evolves from a distribution initially peaked near ϕ ≈ 0 to a bimodal distribution with peaks near ϕ ≈ −1 and ϕ ≈ 1.

Equivalently, we can express the state of the system in terms of a local concentration c(**r**, t) of the liquid ordered phase that can be related to the ϕ(**r**, t) as

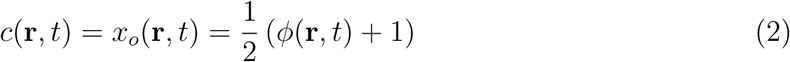

To compute the local lipid compositional order parameter ϕ(**r**, t), we define a two-dimensional plane on both the upper and lower membrane leaflet. In MARTINI simulations, we use C2A, C2B beads for DPPC, D2A, D2B beads for DUPC, and R2 beads for CHOL to define the 2D plane. In atomistic simulations, C27 and C37 atoms define the 2D membrane plane for DPPC and DUPC, while C8 atom defines it for CHOL. These 2D planes are then divided into local areas using either a grid-based definition or Voronoi tessellations.

For the grid-based definition, we divide the total area into NxN grids. The compositional order parameter ϕ(**r**, t) is determined based on the chemical identities of nearest neighbors within the same leaflet by determining the local mole fraction x*_o_*, of DPPC and CHOL, and the local mole fraction x*_d_*, of DUPC.

The effectiveness of the local composition order parameter ϕ to track the progress of the phase separation process, was evaluated by analyzing the probability distributions P(ϕ) at the initial and final phases of the phase separation process for both AA and CG trajectories, as illustrated in Figure 5. ϕ was calculated using an 8 × 8 grid. Initially, the distribution displayed an approximately Gaussian-type distribution centered at zero, indicating the random miscible nature of the lipid mixture. However, as the system evolves towards the final phase-separated stage, the distribution develops high peaks at ϕ ≈ ±1, signifying the formation of L*_o_* and L*_d_* domains.

**Figure 5:**
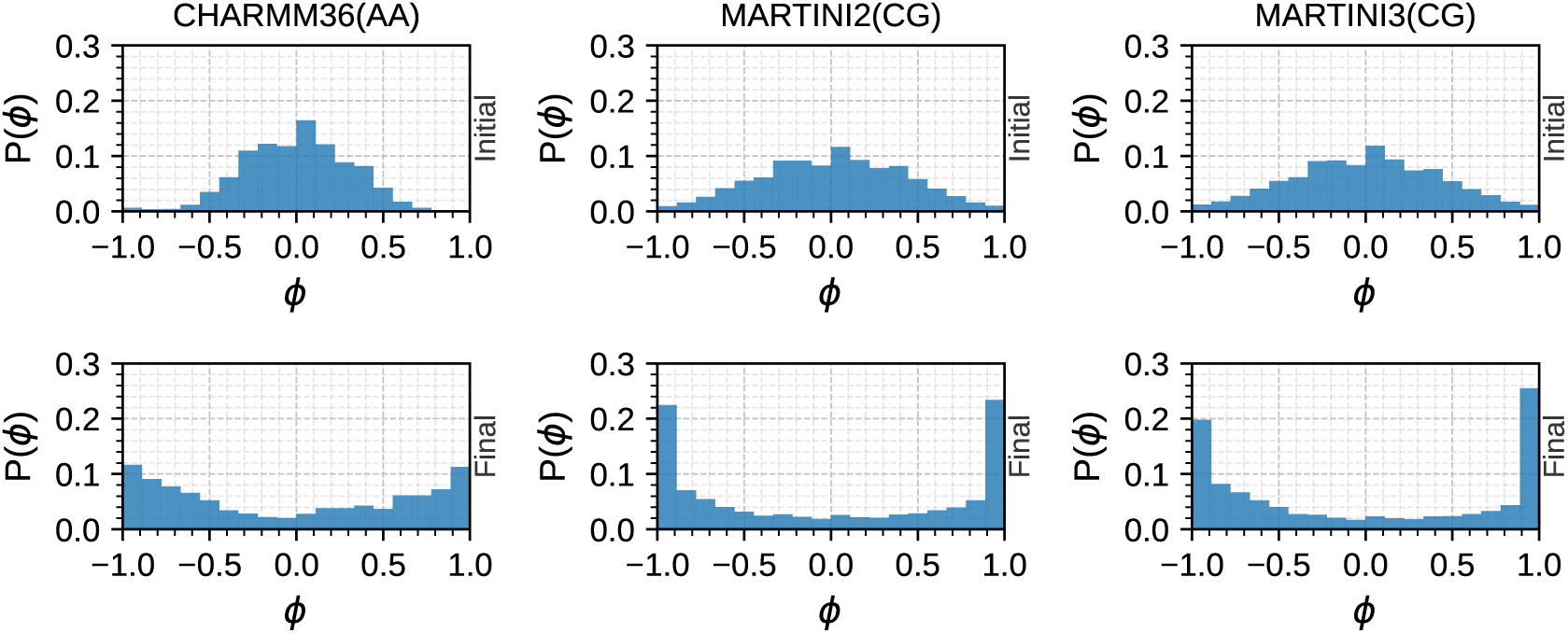
Probability distribution of ϕ at initial and final stages of lipid phase separation for AA(CHARMM36) and CG(MARTINI2 and MARTINI3) simulations. Initial phases demonstrate a gaussian distribution centered at 0 and the final phases exhibit a bimodal distribution with peak at ϕ ≈ ±1.

Figure 6 illustrates the instantaneous values of ϕ(**r**; t) associated with each lipid in the upper leaflet of the membrane as a function of the lipid coordinate calculated at timestamps labeled on the figure. The calculation of ϕ in this case and the succeeding analysis is based on Voronoi Tessellation, as explained above. Initially, at t=0, the different values of ϕ are scattered throughout the plane, and a larger proportion of the plane is occupied with ϕ values near zero, indicating a miscible state. As the phase separation progressed, we observed localization and aggregation of points with similar ϕ values, analogous to that of crystal nucleation. Subsequently, we observed the merger of these small aggregates to form localised larger blue (ϕ ∼ +1) and red (ϕ ∼ −1) domains, signifying the formation of L*_d_* and L*_o_* phases.

**Figure 6:**
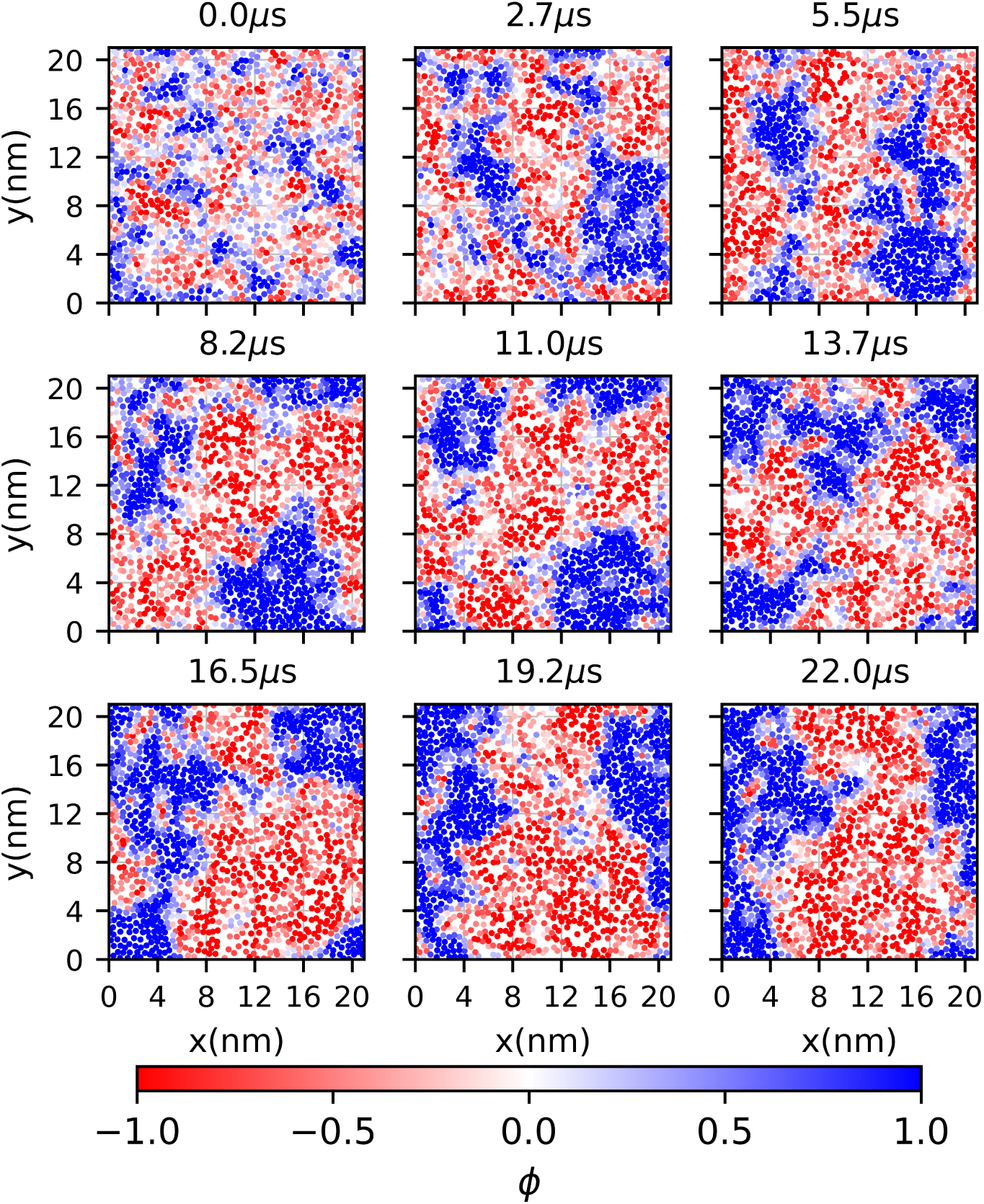
Instantaneous spatial distribution of ϕ as a function of coordinates in XY plane. Each point represent a lipid on the upper leaflet of the membrane. Red colored points represent a negative value of ϕ whereas blue points represent positive values as per the colorbar given below the plots. At t=0, higher as well as lower values of ϕ are observed to be scattered throughout the plane representing a miscible state and as the phase separation progresses higher and lower values of ϕ start to cluster to form larger phase separated domains with blue representing the L*_o_* domain and red representing the L*_d_* domain.

Subsequently, the different stages of the phase separation process is evaluated from the time evolution of the variance of ϕ, ⟨Δϕ^2^⟩. The time evolution of the variance of ϕ(**r**, t) across all points, ⟨Δϕ^2^⟩(t) is defined as

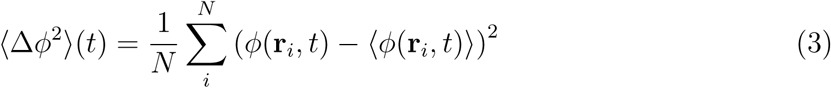

where ϕ(**r***_i_*, t) is the local composition order parameter at a particular position **r***_i_* at time t and ⟨ϕ(**r***_i_*, t)⟩= ϕ(**r***_i_*, t), and N is the number of tesselation cell.

The time evolution of ⟨Δϕ^2^⟩ can be divided into three stages, as shown in Figure 7a:

a. Exponential Stage: This stage is characterized by the rapid exponential growth, ⟨Δϕ^2^⟩ = a_0_e*^α^*^t^ of ⟨Δϕ^2^⟩, where β is the exponential factor and a_0_ is the constant pre-factor of the exponential function, stemming from the initial relaxation of the system leading to formation of small aggregates of similar phases as observed in Figure 6 (Figure 7). This stage lasts until 20 ns in the MARTINI2 timescale. The fitting parameters are provided in Table S1.
b. Power Law Stage: Following the exponential phase, the phase separation slows and transitions to a power-law stage ⟨Δϕ^2^⟩(t) = a_1_t*^β^* that persists up to 3.2 µs in MARTINI2 simulations and 2.8 µs in MARTINI3 simulations (Figure 7). It is characterized by exponents of β*_AA_* = 0.125, β*_M_* _2_ = 0.123, and β*_M_* _3_ = 0.154. In this stage, lipids of similar phases coalesce to form one large L*_o_* or L*_d_* domain, leaving behind nanoscopic impurities in opposing phases. Fitting the cluster size with a power law function, we obtained an exponent of <u>^1^</u>, similar to that from time-dependent spectroscopic analysis of domain growth in near-critical lipid mixtures. ^83,84^ However, as the domain size and ⟨Δϕ^2^⟩ measured here are microscopic quantities, we simply consider these exponents to be measures of the similarity of kinetics between the coarse-grained and all-atom models. The complete set of fitting parameters can be found in Table S2.
c. Plateau Stage: After the formation of larger domains, the phase separation kinetics transitions to a plateau stage as illustrated in the shaded cyan region in Figure 7. This stage has also been observed by Bagchi and co-workers, ^82,85^ who found this logarithmic phase to correspond to the smoothing of domain boundaries. Here, nanoscopic domains of the opposite phase to the major domains gradually merge with the major domain of the corresponding phase.

**Figure 7:**
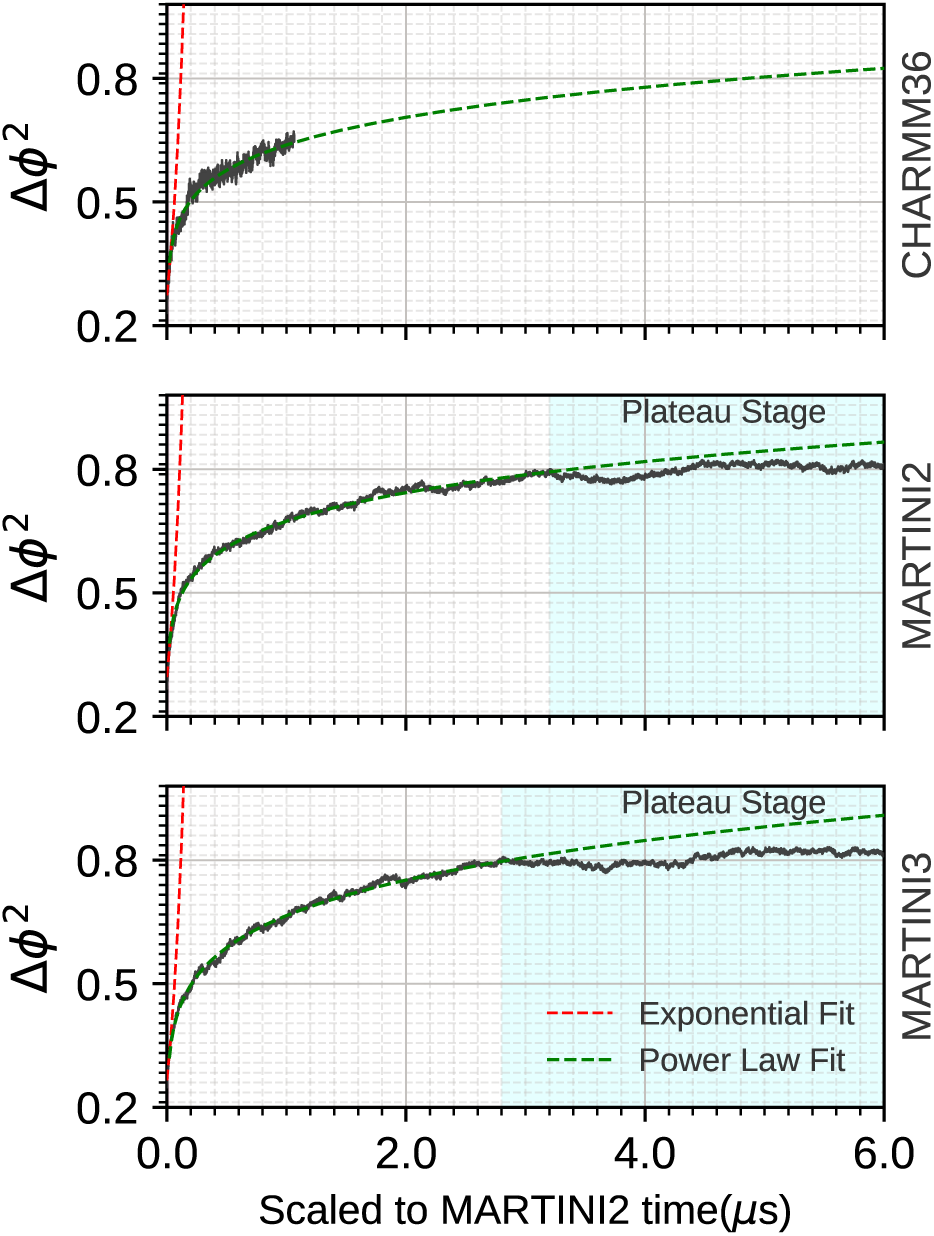
Time evolution of variance of ϕ, Δϕ^2^ averaged over all space with respective fitting curve for exponential phase (red dashed line) and power law phase (green dashed line). In case of MARTINI2 and MARTINI3 simulations the final plateau phase is shaded cyan.

### Higher APP C99 dimer population observed in CG simulations

Figure 8 illustrates the fraction of total simulation frames in which the APP C99 protein is present in dimer state. The dimer state is defined based on the center-of-mass distance between the two proteins: if the center-of-mass distance calculated from the backbone atoms is less than 20 Å, the proteins are classified as a dimer. In CHARMM36 simulations, the protein predominantly remains in the monomer form. In contrast, MARTINI2 and MARTINI3 simulations exhibit a higher population of the protein in the dimer state. Notably, in MARTINI2 simulations, four out of five replicas transition to the monomer form within the first 5 µs as shown in Figure S4. In MARTINI3 simulations, all replicas transition to the monomer form within the first 2.1 µs. The discrepancies between the AA and CG simulations may arise from timescaling. For instance, in CG simulations, some transitions to the dimer state occur within 2.1–5 µs. In our particular case, the timescaling factors for AA simulations relative to MARTINI2 and MARTINI3 are approximately 22 and 29, respectively. This suggests that the 22 µs of sampling in the AA simulations may correspond to only about 0.7-1.0 µs of effective CG timescales, potentially making it insufficient for the protein to achieve the dimer state.

**Figure 8:**
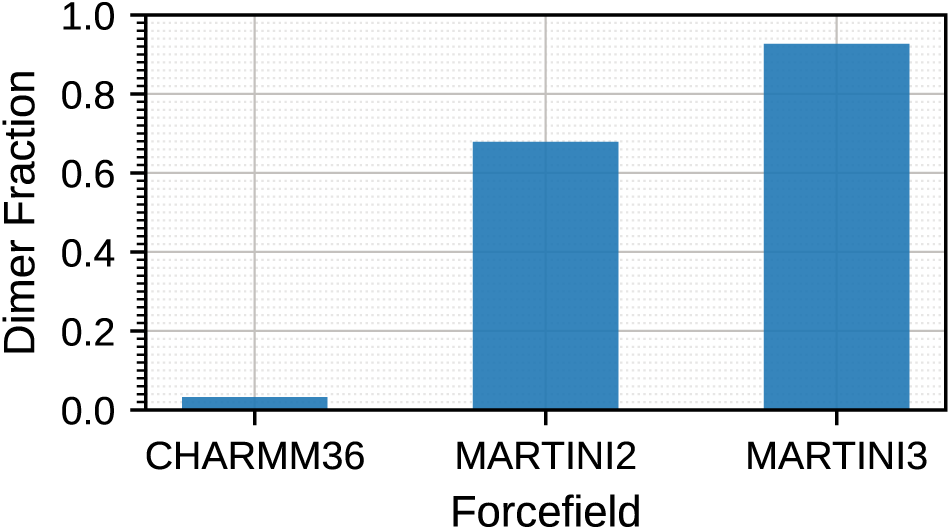
Population of APP C99 dimer observed in AA (CHARMM36) and CG (MARTINI2 and MARTINI3) simulations. The dimer population was calculated based on the center-of-mass distance between the backbones of two APP C99 monomers. A center-of-mass distance less than 20 Åwas classified as a dimer.

### Similar ordering but distinct composition of lipids surrounding the protein

Differences in protein-lipid interaction parameters across force fields are known to influence the local environment of proteins within membranes. ^86–90^ In complex bilayers, two key aspects can be affected: the local lipid composition surrounding the protein and the ordering of lipids around it. To investigate lipid ordering, we calculated the |Ψ_6_| parameter for lipids in the first Voronoi shells around the protein. The results, shown in Figure 9, indicate similar |Ψ_6_| distributions for both all-atom (AA) and coarse-grained (CG) force fields, with peaks in the range of |Ψ_6_| 0.2–0.4. These low |Ψ_6_| values suggest a disordered lipid packing around the protein, which is consistent with its localization in the L*_d_* domain of the membrane.

**Figure 9:**
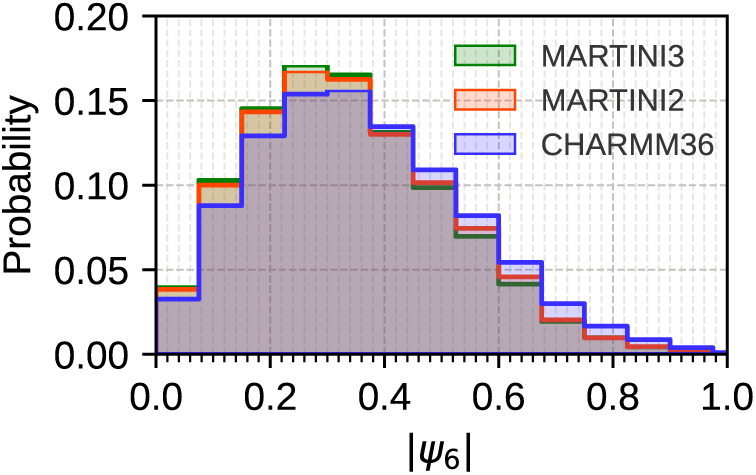
Probability distribution of Ψ_6_ values of first voronoi neighbours around the protein.

In contrast, the lipid composition surrounding the protein differs significantly for varying force fields as shown in Figure 10. In CHARMM36 and MARTINI3 simulations, the protein is predominantly surrounded by DUPC, with peak fractions of 0.8 and 1.0, respectively. However, in MARTINI2 simulations, while DUPC is still the major lipid, substantial proportions of DPPC and CHOL are also present around the protein. This compositional difference stems from the protein’s preferential localization within the membrane. In MARTINI2, the protein is localized in the L*_o_*-L*_d_* domain interface, an often-observed phenomenon in the MARTINI2 force field. ^47,91^ In MARTINI3 and CHARMM36, C99 is more embedded within the bulk L*_d_* domain, in line with the ∼90% preference for C99 to partition to L*_d_* domains. ^20,23^

**Figure 10:**
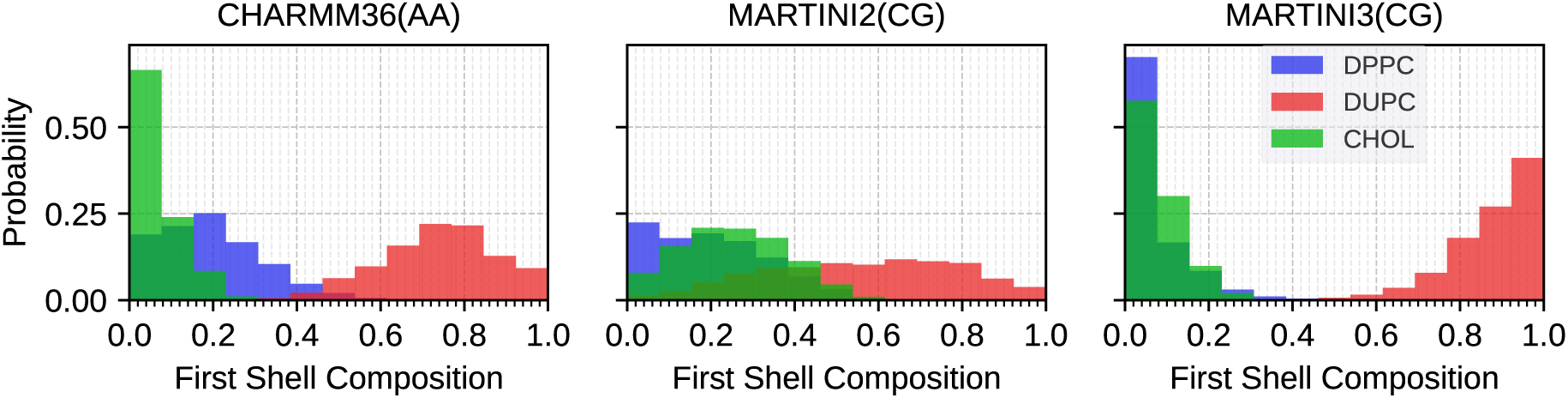
Probability distribution of DPPC (blue), DUPC(red) and CHOL(green) in the first voronoi neighbouring shell of APP C99 for AA (CHARMM36) and CG (MARTINI2 and MARTINI3) force fields.

Despite differences in protein co-localization and oligomeric states, the mechanism and kinetics of phase separation in the presence of the protein are not affected. This suggests that while local lipid composition, protein organization, and lipid-protein interactions vary between force fields, the larger-scale phase behavior of the membrane remains robust. The thermodynamic driving forces for phase separation appear to dominate over the local variations in protein environment, as well as the differences in lipid-lipid interactions and lipid diffusion, establishing consistent phase separation kinetics across different scale of simulations.

## Conclusions

In this work, molecular dynamics simulations of *de novo* phase separation in ternary lipid mixtures consisting of 37% DPPC, 37% DUPC, and 26% cholesterol are performed for both AA (CHARMM36) and CG (MARTINI2 and MARTINI3) models. By analyzing the dynamics of lipid phase separation from a miscible state to a distinctly phase separated state, we have gained insight into the time scale and mechanism of lipid phase separation. Comparison of the dynamics for the AA and CG models leads to insight into the origin of the differences in domain formation rate observed for the two models.

The time scale for phase separation in AA models is observed to be on the order of 20 µs, while the time scale for phase separation in simulations based on CG models is found to be faster by a factor of 22 for MARTINI2 simulation and 29 for MARTINI3 simulations. The faster rate of domain formation is partly explained by an enhanced rate of diffusion in CG as opposed to AA models. However, an additional difference is observed in the greater lipid-lipid sticking probability, the fraction of monomer collisions with a cluster that result in a monomer joining the cluster, for CG models in the process of nucleation and growth of domains.

A three-stage mechanism was observed for lipid phase separation in AA and CG models. At short times, we observe an exponential growth leading to the formation of nanoscopic liquid ordered domains. These small domains subsequently merge to form larger domains, and this stage is characterized by power law growth. At long times, we observe a plateau stage. Bagchi and coworkers, ^82,85^ also observed a similar plateau phase in binary liquid phase separation which they attributed to logarithmic dynamics. This study provides insight into the nature of lipid phase separation and the formation of nanodomains in lipid mixtures.

Notably, protein-lipid interactions exhibited force-field-dependent variations that influenced the local lipid environment around the protein as expected. Despite these local differences, the larger-scale phase behavior of the membrane, including the mechanism and kinetics of phase separation, remained robust across models. These same regimes of kinetic behavior may be reproducible in more aggressively coarse-grained models. ^92^ Taken together, our results provide insight into the time scale and mechanism of phase separation in ternary lipid mixtures in presence of a transmembrane protein. The findings of this study provide insight into the mechanism of nanodomain formation relevant to our understanding of the size and time scale of transient nanodomain formation in membrane.

## Acknowledgement

The authors gratefully acknowledge the generous support of the National Science Foundation (CHE-1900416), the National Institutes of Health (R01 GM107703), and the highperformance computing resources of the Boston University Shared Computing Cluster (SCC). The authors thank Ayan Majumder, Biman Bagchi and Sarmistha Sarkar for helpful discussions.

