## Supplementary Information for "Exploring the kinetics and mechanism of phase separation in ternary lipid mixtures containing APP C99 using atomistic vs coarse-grained MD simulations"

### Modeling nucleation-growth kinetics

We can model the rate of nucleation and growth of a growing cluster if $i$ as

$$\mathrm{n(i)} \underset{e(i+1)}{\overset{c(i)}{\rightleftharpoons}} \mathrm{n}(i+1)$$

where $c(i)$ is the rate of condensation of a monomer onto a cluster of $i$ monomers and $e(i+1)$ is the rate of evaporation of a monomer from a cluster of $i+1$ monomers. If we ignore evaporation, the kinetics feeds forward toward the formation of larger and larger clusters.

For gas phase dynamics, we imagine particles moving ballistically through space with a thermal distribution of velocities. The probability of colliding with a cluster will depend on the collisional cross section of the cluster $4\pi r_i^2$ where $r_i$ is the radius of a cluster of $i$ monomers. We approximate the rate

$$c(i) = \alpha\, 4\pi r_i^2 \sqrt{\frac{kT}{2\pi m_1}}\, n_1(t)$$

which is also commonly expressed as

$$c(i) = \alpha\, 4\pi r_1^2 \sqrt{\frac{kT}{2\pi m_1}}\, i^{\frac{2}{3}} n_1(t)$$

where $r_i = r_1 i^{\frac{2}{3}}$ The parameter $\alpha$ is a sticking probability that represents the fraction $0 \le \alpha \le 1$ of collisions that lead to the addition of a monomer to the cluster.

Alternatively, for diffusion controlled reactions the rate of encounter of the diffusing monomer and a growing cluster of radius $r_i$ is

$$c(i) = \alpha\, 4\pi D\sigma\, n_1(t)$$

where $D = D_i + D_1$ is the relative rate of diffusion of the cluster and monomer and $\sigma = r_i + r_1$

is the encounter radius for contact between the cluster and monomer. As in the case of the gas phase collisional rates, $\alpha$ is a sticking probability. For the case of nucleation and growth of nanodomains in a lipid membrane we assume that encounters between diffusing monomers and clusters is diffusion controlled.

We note that following classical homogeneous nucleation theory, the nucleation rate is defined

$$J = \left[ \frac{1}{c(1)n_1} + \sum_{i=2}^{\infty} \frac{1}{c(i)n_{\rm eq}(i)} \right]^{-1}$$

where $n_{\rm eq}(i)$ is the equilibrium number density of clusters composed of $j$ monomers and $c(i)$ is the rate of at which a cluster of size $j$ acquires an additional monomer to become a cluster of size $i+1$. However, this result is only relevant when the distribution of cluster sizes conforms to the equilibrium distribution, which is not appropriate in a non-equilibrium dynamics of cluster growth most appropriate to our lipid phase separation dynamics.

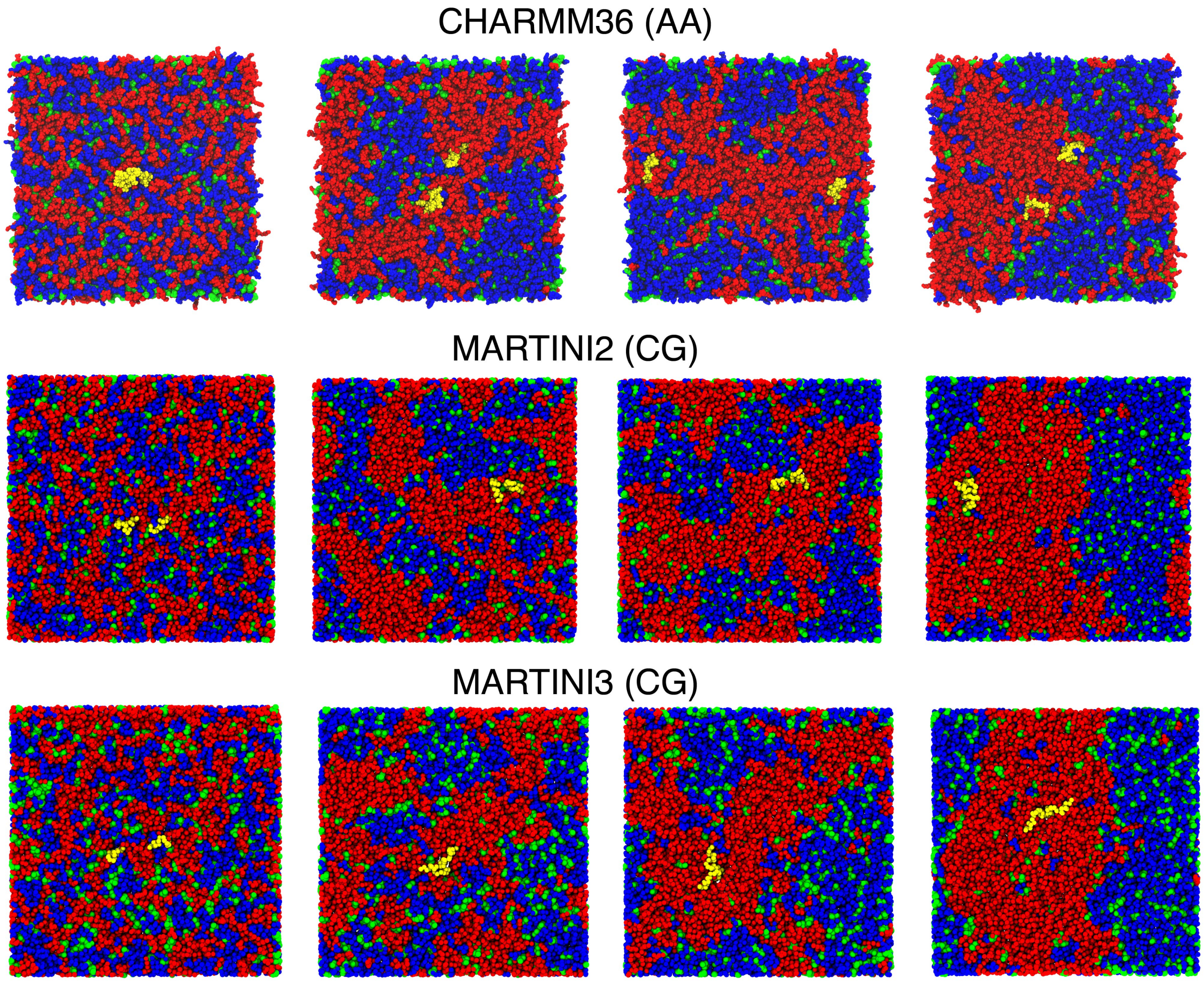


Figure S1: Four instantaneous configurations depicting the process of lipid phase separation of DPPC (blue), DUPC (red), and cholesterol (green) into $L_o$ (blue) and $L_d$ (red) regions for the AA(CHARMM36m) trajectory and a CG(MARTINI2 and MARTINI3) trajectory. The extent of phase separation is comparable in the AA and CG models.

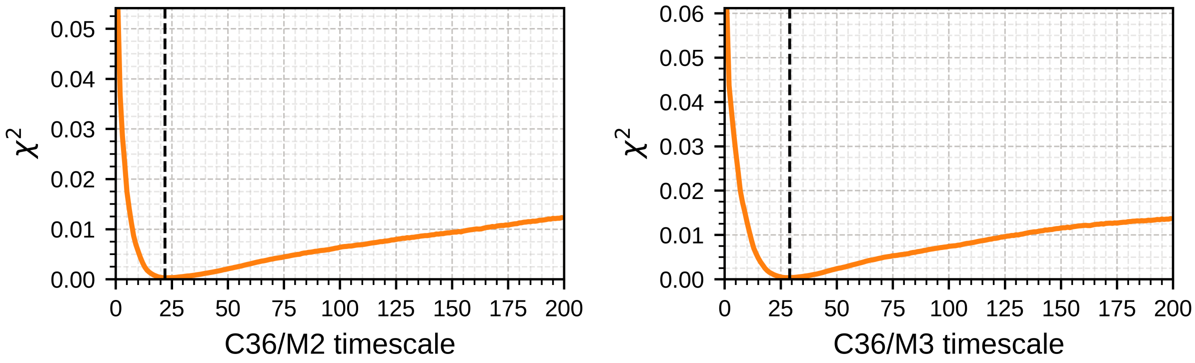


Figure S2: $\chi^2$ plotted as a function of timescaling between AA (CHARMM36m) and (a) MARTINI2(M2) (b) MARTINI3(M3). For CHARMM36 and MARTINI2 simulations the scaling factor was found to be 22 and for CHARMM36 and MARTINI3 simulations it was found to be 29 based on minimum $\chi^2$ value.

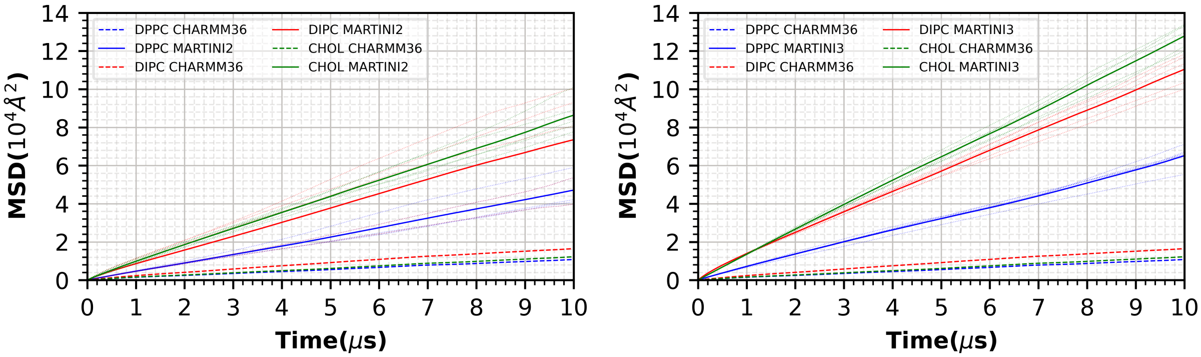


Figure S3: Mean square displacement of DPPC, DUPC and Cholesterol with CHARMM36m, MARTINI2 and MARTINI3 models

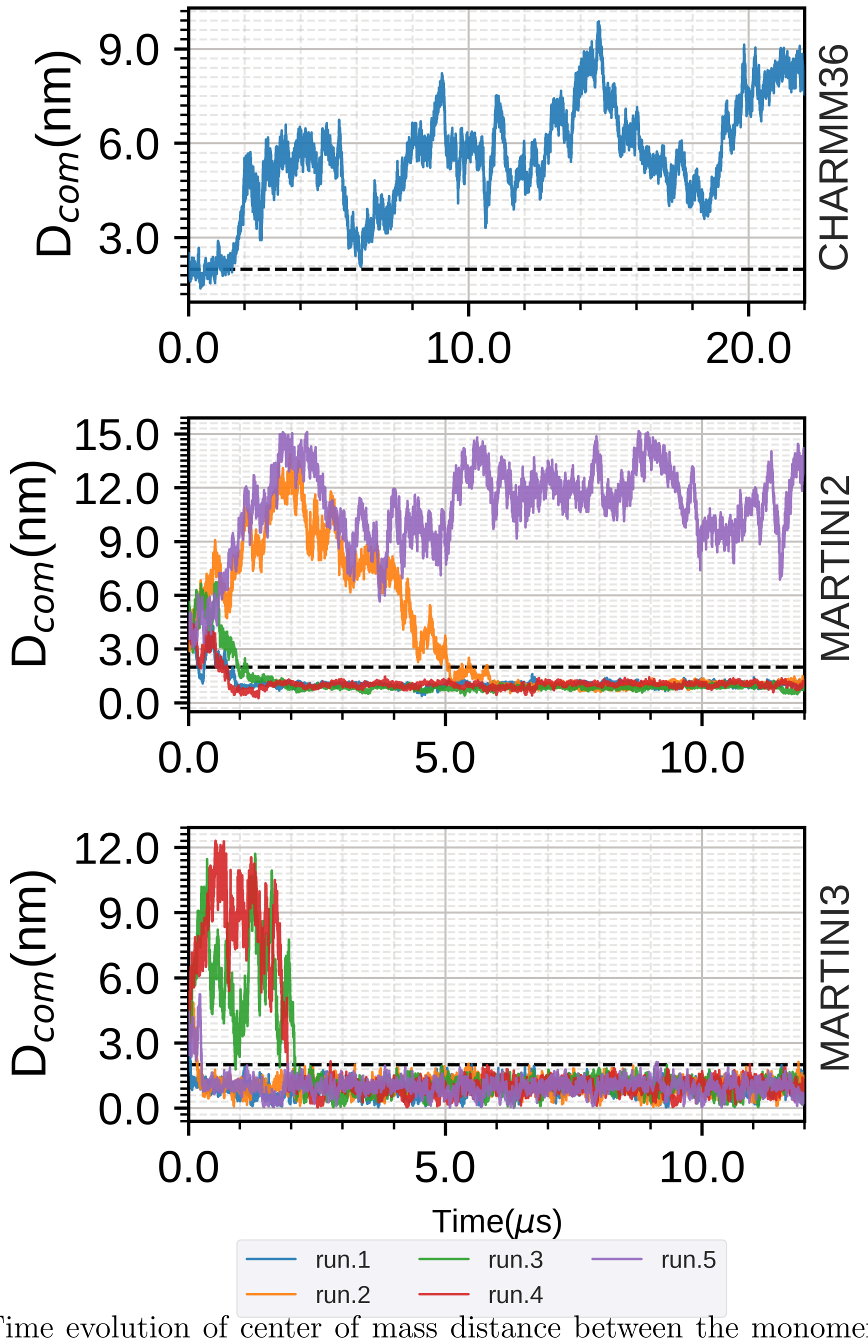


Figure S4: Time evolution of center of mass distance between the monomers of APP C99 protein in AA (CHARMM36) and CG (MARTINI2 and MARTINI3) forcefields. The time evolution of 5 independent production run are shown in the MARTINI2 and MARTINI3 panels.

Table 1: Fitting parameters of the exponential phase. Linear regression based on $\ln \langle \Delta\phi^2 \rangle = \alpha\mathrm{t} + \ln a_0$ provides the fit.

| Force-field | $\alpha$ | $\ln a_0$ |
|---|---|---|
| AA (CHARMM36) | $7.57 \pm 0.104$ | $-1.17 \pm 0.001$ |
| MARTINI2 | $7.66 \pm 0.266$ | $-1.11 \pm 0.003$ |
| MARTINI3 | $7.74 \pm 0.289$ | $-1.18 \pm 0.003$ |

Table 2: Fitting parameters of the power law phase. Linear regression based on $\ln \langle \Delta\phi^2 \rangle = \beta \ln t + \ln a_1$ provides the fit.

| Force-field | $\beta$ | $\ln a_1$ |
|---|---|---|
| AA (CHARMM36) | $0.125 \pm 8.6\text{x}10^{-5}$ | $-0.48 \pm 0.0001$ |
| MARTINI2 | $0.123 \pm 0.0001$ | $-0.438 \pm 0.0001$ |
| MARTINI3 | $0.154 \pm 0.0001$ | $-0.449 \pm 0.0001$ |
